# Effects of dietary taurine level on visual function in European sea bass (*Dicentrarchus labrax*)

**DOI:** 10.1101/577023

**Authors:** Richard W. Brill, Andrij Z. Horodysky, Allen R. Place, Mary E.M. Larkin, Renate Reimschuessel

**Author notes:** Corresponding author (RB).

## Abstract

Dietary insufficiencies have been well documented to decrease growth rates and survival (and therefore overall production) in fish aquaculture. By contrast, the effects of dietary insufficiencies on the sensory biology of cultured fish remains largely unstudied. Diets based solely on plant protein sources could have advantages over fish-based diets, because of the cost and ecological effects of the latter, but lack the amino acid taurine. Adequate levels of taurine are, however, necessary for the development of a fully functional visual system in mammals. As part of ongoing studies to determine the suitability of plant-based diets, we investigated the effects of normal and reduced taurine dietary levels on retinal anatomy and function in European sea bass (*Dicentrarchus labrax*). We could not demonstrate any effects of dietary taurine level on retinal anatomy, nor the functional properties of luminous sensitivity or temporal resolution (measured as flicker fusion frequency). We did, however, find an effect on spectral sensitivity. The peak of spectral sensitivity of individuals fed a 5% taurine diet was rightward shifted (i.e., towards longer wavelengths) relative to that of fish fed a 0% or 1.5 % taurine diet. This difference in in spectral sensitivity was due to a relatively lower level of middle wavelength pigment (maximum absorbance ≈500 nm) in fish fed a 5% taurine diet. Changes in spectral sensitivity resulting from diets containing different taurine levels are unlikely to be detrimental to fish destined for market but could be in fishes that are being reared for stock enhancement programs.

## Introduction

Carnivorous fishes cannot synthesize many essential amino and fatty acids and must receive them through their diet [1,2]. Consequently, the aquaculture of these species requires diets based on, or at least supplemented with, protein sources from wild-caught fish. Up to two to five times more fish is, however, required to culture a product than is provided by it [3]. This increases the cost of production and can have ecological consequences as capture of wild-caught fish may impact forage fish populations and the higher trophic level species that feed on them [4,5]. An equally important concern is the potential for fishmeal to contain xenobiotic compounds such as polychlorinated biphenyls (PCB’s) and mercury. These undergo biomagnification leading to elevated levels of contaminates in the endproduct. In brief, fishmeal diets for the aquaculture of carnivorous fishes are costly, and potentially unsustainable as well as unsafe. Purely plant-based diets circumvent these issues, but generally lack essential and semi-essential amino acids (e.g., taurine, methionine, lysine), as well as many vitamins and minerals needed in microquantities [6]. Plant-based diets are therefore currently supplemented with fishmeal or fish oil [7,8,9]. But we argue, as have others [10)], that there remains an exigent need to determine if purely plant-based diets can support survival rates, growth rates, and feeding efficiencies (i.e., the ratio of the mass of fish produced per mass of feed) necessary for the successful aquaculture of carnivorous fishes.

Omnivorous species have been the easiest to convert to low or no fishmeal diets; whereas marine carnivorous species have been the most difficult [11]. But it is the latter that are produced by aquaculture for restocking programs (i.e., to augment wild populations [12]). Diets differing in fatty acid composition can have significant impacts on the growth performance, energetics, cardiorespiratory physiology, hypoxia tolerances, and exercise and recovery performance of fishes [13–17]. It is therefore plausible that the metabolic performance and hypoxia tolerances of fishes fed plant-based diets may be significantly different – with major implications for the suitability of fishes for re-stocking programs.

More specific to our project, plant protein sources lack taurine. Taurine is a sulfur containing amino acid that is found in higher concentrations than any other free amino acid (i.e., amino acids not incorporated into any known proteins) and its roles in the proper development and function in a variety of vertebrate tissues have received considerable attention [18–21]. Taurine is considered a conditionally indispensable amino acid for humans and non-human primates and an essential amino acid in some mammalian carnivores (e.g., felines) [20], but little attention has been paid to the required levels of this amino acid or its roles in fishes. We hypothesize that most marine carnivorous fishes lack the little ability to synthesize taurine (due to the large quantities found in their natural prey items) and therefore require it to be supplied in the diet. We also hypothesize that plant-based diets may not allow for development of fully functional visual system as a low taurine diet, or treatments with agonists of taurine uptake, have been documented to impede the embryonic development of the retina and maintenance of normal retinal function in mammals; the latter because of taurine’s role as an antioxidant and osmolyte (i.e., a compound maintaining intracellular osmotic balance) [20–24]. High concentrations of taurine have, moreover, been found in the photoreceptor cells (i.e., rod and cone cells) and retinal pigment epithelium in several teleost fish species [25–30]. Although it has been suggested that taurine is only an osmolyte in retinal cells of fishes [26], other investigators have concluded that the role of taurine in development and maintenance of retinal function is conserved throughout the vertebrate order [21,31]. We therefore posit that diets containing inadequate levels of taurine could result in diminished visual system function in carnivorous fishes. We recognize that less than fully functional visual systems are unlikely to be detrimental to fish destined for market, yet we contend that a diminished functionality of the visual system of fishes being cultured for stock enhancement programs would decrease their survival and fitness (e.g., growth and reproduction) relative to wild individuals. If this is the case, programs rearing fish for restocking would be less able to meet their ultimate objective. Therefore, in conjunction with our ongoing study examining the overall efficacy of formulations of our plant-based diet for generating acceptable growth rates in European sea bass, we expanded our efforts to examine the effects of dietary taurine level on visual function.

## Materials and Methods

Our study was carried out under protocols approved by the Institutional Animal Care and Use Committees of the University of Maryland Baltimore Medical School and the College of William and Mary and followed all applicable laws and regulations. European sea bass were obtained from the Aquaculture Research Center at the Institute of Marine and Environmental Technology (IMET, Baltimore, MD). The average starting weight was ~15 g for fish subsequently reared on the 5% taurine diet and ~25 g for fish subsequently reared on the 0 or 1.5% taurine diet. Fish were divided by diet and housed in eight-foot diameter, four cubic meter recirculating systems with shared mechanical and life support systems. The latter included a protein skimmer, ozonation, mechanical filtration (in the form of bubble-bead filters), and biological filtration. Water quality (measured two to three times per week) was not significantly different between systems (ANOVA, p>0.05). Mean (± SEM) water quality values in the tanks were: dissolved oxygen 5.7 ± 1.6 mg L^-1^, temperature 27 ± 2 ^o^C, pH 7.6 ± 0.3, total ammonia nitrogen (NH_3_) 0.06 ± 0.06 mg L^-1^, nitrite (NO_2_^-^) 0.12 ± 0.08 mg L^-1^, nitrate (NO_3_^-^ ) 49 ± 9 mg L^-1^, alkalinity 96 ± 23 meq L^-1^, and salinity 25 ± 2 ppt. Fish were fed 3.5% of their body weight per day and maintained on each specific diets for five to six months.

### Diet preparation

The three diets formulations (Table 1) were prepared by Zeigler Bros. (Gardners, PA, USA) and analysis of their proximate composition (Table 2) was performed by New Jersey Feed Laboratory, Inc. (Ewing Township, NJ, USA).

**Table 1.** Formulations for the three experimental diets.

| Constituent | 0% taurine | 1.5% taurine | 5% taurine |
| --- | --- | --- | --- |
| Profine VF | 28.75 | 28.75 | 28.75 |
| Soybean meal, 47.5% | 23.33 | 23.33 | 23.33 |
| Wheat flour, bagged | 16.54 | 15.04 | 11.54 |
| Corn gluten, 60% | 15.34 | 15.34 | 15.34 |
| Menhaden gold oil, top-dressed | 5.96 | 5.96 | 5.96 |
| Monocalcium phosphate FG | 3.95 | 3.95 | 3.95 |
| Lecithin FG | 3 | 3 | 3 |
| L-Lysine, 98.5% | 0.75 | 0.75 | 0.75 |
| Choline chloride, 70% | 0.6 | 0.6 | 0.6 |
| Potassium chloride FG | 0.56 | 0.56 | 0.56 |
| DL-Methionine, 99% | 0.45 | 0.45 | 0.45 |
| Sodium chloride | 0.28 | 0.28 | 0.28 |
| Vitamin C | 0.2 | 0.2 | 0.2 |
| Premix AquaVit | 0.12 | 0.12 | 0.12 |
| Premix Aquamin Fish | 0.12 | 0.12 | 0.12 |
| Magnesium oxide FG | 0.05 | 0.05 | 0.05 |
| Taurine FG | 0 | 1.5 | 5 |

FG = food grade

**Table 2.** Proximate composition (%) of three experimental diets.

| Constituent | 0% taurine | 1.5% taurine | 5% taurine |
| --- | --- | --- | --- |
| Moisture | 8.05 | 6.97 | 9.5 |
| Protein | 43.82 | 45.02 | 46.65 |
| Fat | 5.94 | 6.19 | 6.94 |
| Fiber | 3.06 | 1.95 | 2.25 |
| Ash | 7.08 | 7.75 | 7.71 |
| Taurine | 0.07 | 1.48 | 4.85 |

### Tissue sampling for analysis retinal anatomy

Eyes were harvested from fish not used in the ERG experiments. Food was withheld for 24 hours and 10-12 fish from each diet were anesthetized in a bath containing 25 mg L^-1^ MS-222 (Syndel, Ferndale, WA, USA) buffered with 50 mg L^-1^ sodium bicarbonate (Sigma-Aldrich, St. Louis, MO, USA). Upon removal from the anesthesia bath, the spinal cord was immediately severed, and eyes harvested. Retinal tissue was subsequently embedded, sectioned, mounted on glass slides, and stained with Hematoxylin-Eosin using standard histological procedures. Images were captured using a BX53 microscope outfitted with a DP73 camera and visualized with the cellsSens software (version 510) (Olympus, Center Valley, PA, USA).

### Retinal responses to light stimuli

Fish were transferred to the Virginia Institute of Marine Science (Gloucester Point, VA) where whole-animal corneal electroretinography (ERG) was used to assess three standard metrics of retinal function: (1) luminous sensitivity (V log I response), (2) flicker fusion frequency (FFF, a measure of temporal resolution), and (3) spectral sensitivity (32-36). Anesthesia and handling methodologies were as described previously (33-35). Teflon-coated, silver – silver chloride, 0.5mm wire electrodes were used to measure ERG potentials: the active electrode was placed on the corneal surface and a reference electrode in the nasal cavity. All subjects were dark-adapted for a minimum of 60 minutes prior to visual trials. Electrode placements, as well as any further modifications to the experimental setup, were conducted under a dim red LED light source (peak wavelength of 660 nm) that is beyond the spectral sensitivity of European sea bass.

Luminous sensitivity was assessed using stimulus intensities covering six orders of magnitude using a collimated white LED source and neutral density filters progressing from subthreshold to saturation intensity levels in 0.2 log unit steps. FFF was assessed by measuring the ability of the retinal responses to track sinusoidally modulated white light stimuli ranging in frequency from 1 Hz (0 log units) to 100 Hz (2.0 log units), presented in increments of 0.2 log unit frequency steps. FFF was measured at stimulus intensities of 25%, 50% and 100% of the maximum response, as well as fixed light levels (log I = 1.9, 2.7, and 3.7; with I in units of candela per m^2^). Spectral sensitivity was assessed using stimuli over wavelengths from the ultraviolet (300 nm) to the near infrared (700 nm) presented sequentially in 10 nm steps. Monochromatic light flashes (50% bandwidth = 5 nm) were made approximately equally quantal through a series of neutral density filters and subsequently corrected to predict isoquantal responses, as described previously (33-36). To form hypotheses regarding the number and spectral distribution of visual pigments present, and the effects of dietary taurine levels on the distribution of pigments contributing to spectral sensitivity, we fitted the SSH [37] and GFRKD [38] vitamin A1 rhodopsin absorbance templates separately to the photopic spectral sensitivity data. Estimates of the unknown model parameters (λ_max_ values and their respective weighting proportions) were derived by fitting the summed curves to the ERG data using maximum likelihood. We objectively selected the appropriate template (SSH or GFRKD) and number of contributing pigments using an information theoretic approach following Akaike’s information criterion that is a parsimonious measure that strikes a balance between model simplicity and complex overparameterization (39). All parameter optimization, template fitting and model selection was conducted using the software package R version 3.2.2 (R Development Core Team).

Statistical tests comparing the effects of dietary taurine levels on the magnitude of responses to various light levels were done using Sigmaplot (version 11.2, Systat Software, San Jose, CA). Analysis of data were performed using a one-way analysis of variance (ANOVA) test on means when the data were normally distributed, and a Kruskal-Wallis one-way analysis of variance on ranks when the data were not. Comparisons the magnitude of responses to various light levels (i.e., luminous sensitivity) tests were limited to the effects of diet within a given light levels. Statistical tests comparing the effects of dietary taurine levels on FFF were preformed using the two-way repeated measures ANOVA procedure in Sigmaplot, with the Holm-Sidak method to conduct all pairwise multiple comparisons.

## Results and discussion

### Retinal morphology

There were no obvious effects of dietary taurine level on the morphology of identifiable layers in European sea bass retina (Fig 1) not retinal cell layer thickness ratios (Fig 2) equivalent to the massive disruption seen in retinal tissue of domestic cats fed low taurine diets [40].

**Fig 1.**
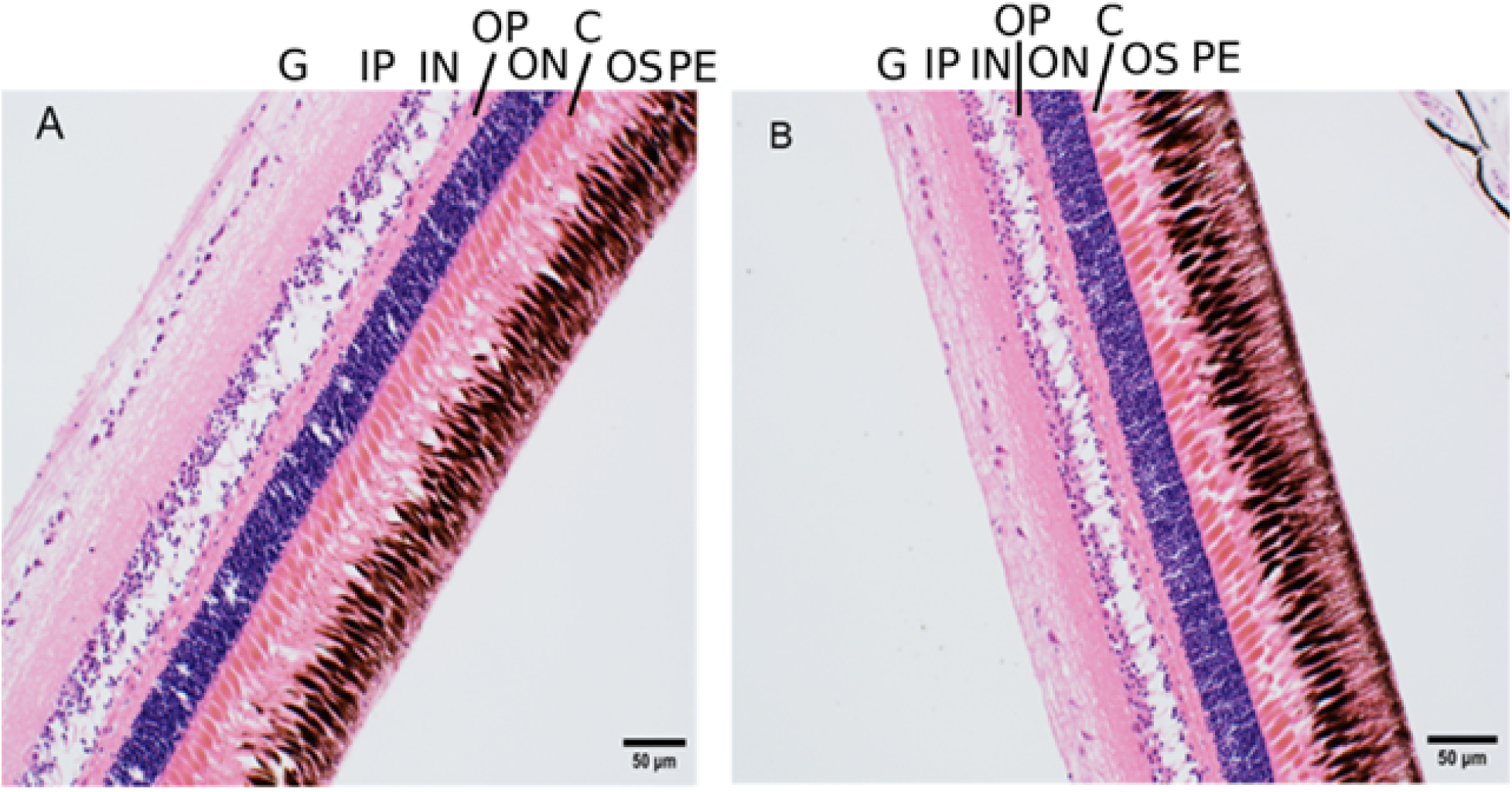
Identifiable layers in European sea bass retinas. Panel A is a representative retina from fish fed a 0% taurine diet, panel B is a representative retina from fish fed a 5% taurine diet. G=ganglion cell layer, IP=inner plexiform layer, IN=inner nuclear layer, OP=outer plexiform layer; ON=outer nuclear layer, C=cone photoreceptors, OS=outer segments of the photoreceptor layer, PE=pigmented epithelium.

**Fig 2.**
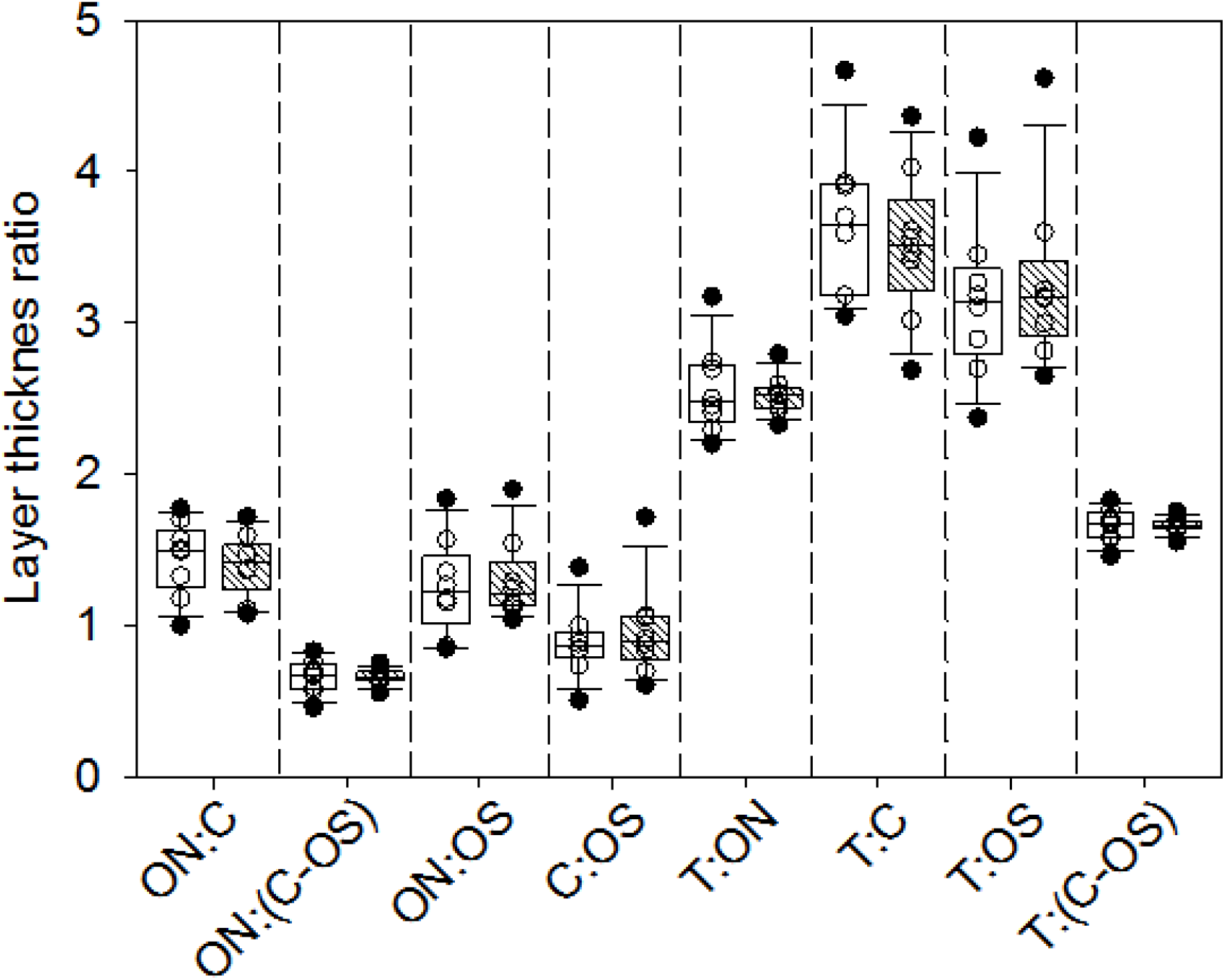
Retinal cell layer thickness ratios in European sea bass fed diets with 0 or 5% taurine (open and cross hatched boxes, respectively). The boundary of the box closest to zero indicates the 25^th^ percentile, the line within the box marks the median, and the boundary of the box farthest from zero indicates the 75^th^ percentile. Whiskers (error bars) below and above the box indicate the 10^th^ and 90^th^ percentiles, respectively. Data points above and below the whiskers are considered outliers. T= total retina thickness, C= cone photoreceptors layer thickness, ON=outer nuclear layer thickness, OS= thickness of the outer segments of photoreceptor layer, IN=inner nuclear layer thickness (μm).

### Retinal function

Retinal responses to increasing light levels showed the expected steep increases up to those levels producing maximum response (Fig 3). When light levels are expressed in log units, retinal response curves were the expected sigmoidal shape (Fig 3 insert). There was only one significant effect of dietary taurine levels on luminous sensitivity in European sea bass, and only at light levels needed to produce a response 75% of maximum (Fig 4). In other words, significantly higher light levels were required to achieve responses above approximately 50% of maximum in fish fed a diet lacking taurine. Flicker fusion frequencies generally showed the expected increases with increasing light intensities, and dietary taurine level had no influence on FFF when comparisons are made at the same light level (Fig 5).

**Fig 3.**
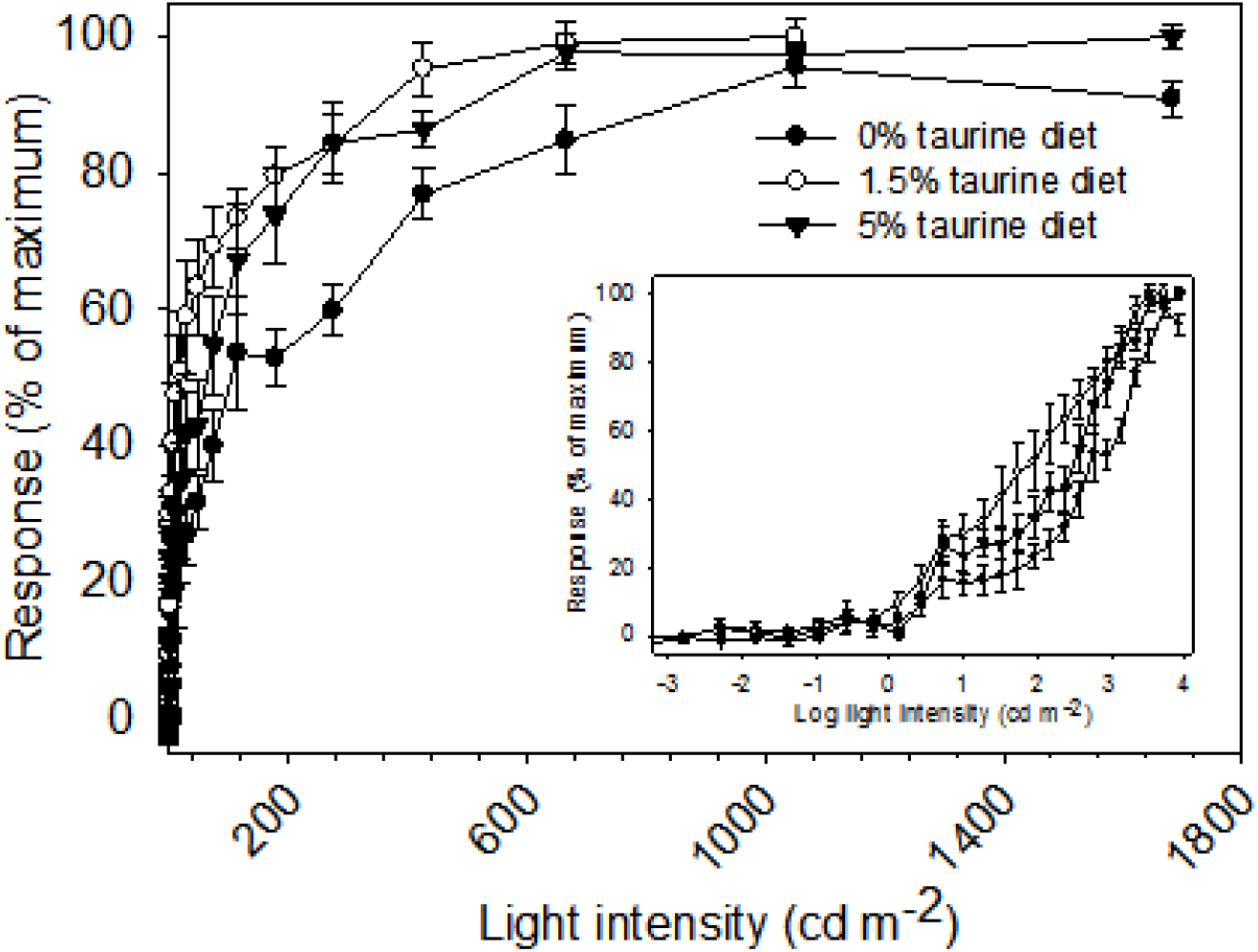
Luminous sensitivity (i.e., intensity–response curves) of European sea bass. Data are mean values (± SEM). Response values for individuals were normalized to 0-100%, the values averaged, and the mean values rescaled to 0-100%. The inset shows the same data, but with light intensities expressed in log units. Increases in light intensity were in 0.2 log unit steps.

**Fig 4.**
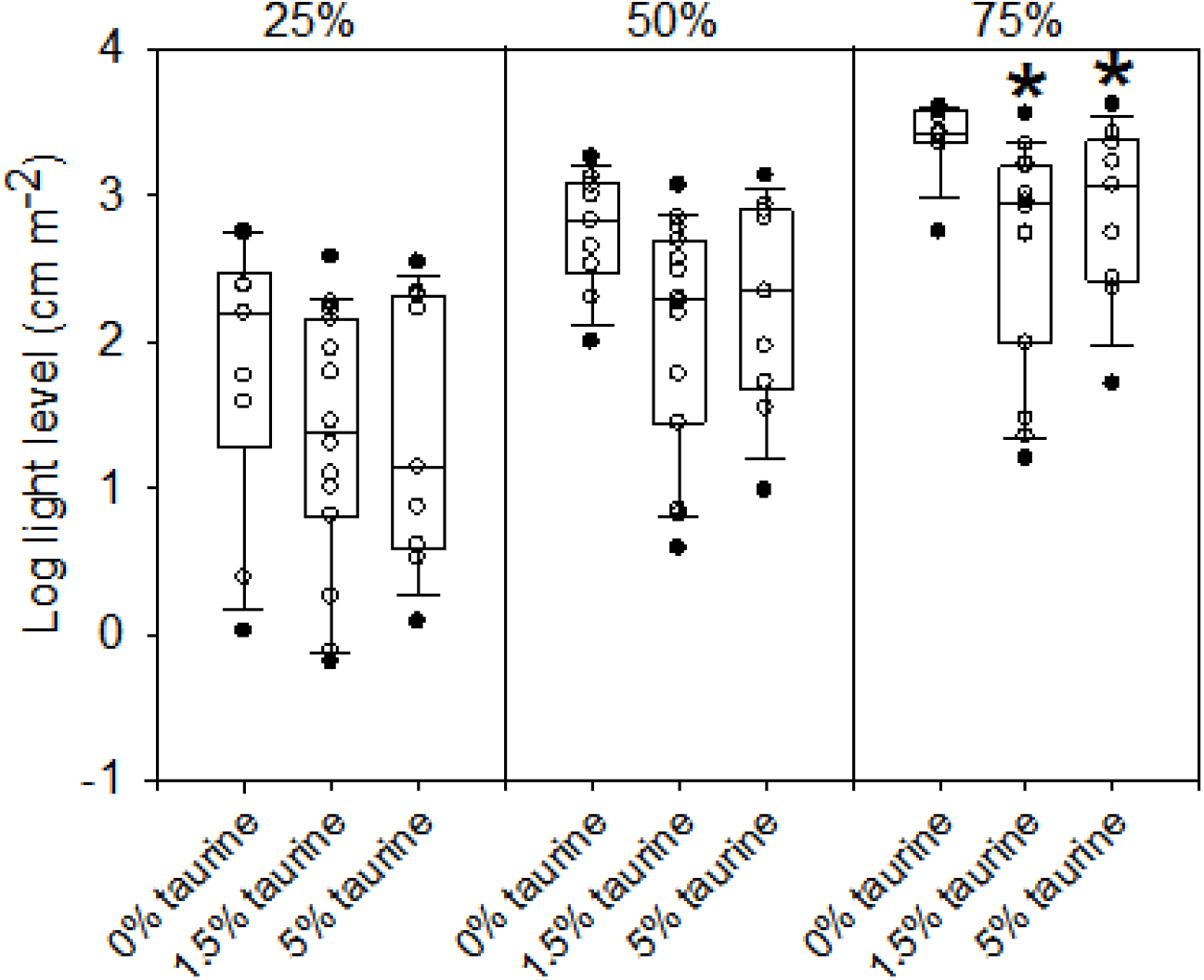
Light levels needed to produce a given response relative to that of the maximin response in intensity–response curves (Fig 3). The boundary of the box closest to zero indicates the 25^th^ percentile, the line within the box marks the median, and the boundary of the box farthest from zero indicates the 75^th^ percentile. Whiskers (error bars) above and below the box indicate the 90^th^ and 10^th^ percentiles. Data points above and below the 90^th^ and 10^th^ percentiles are considered outliers and are not used to determine median values.

**Fig 5.**
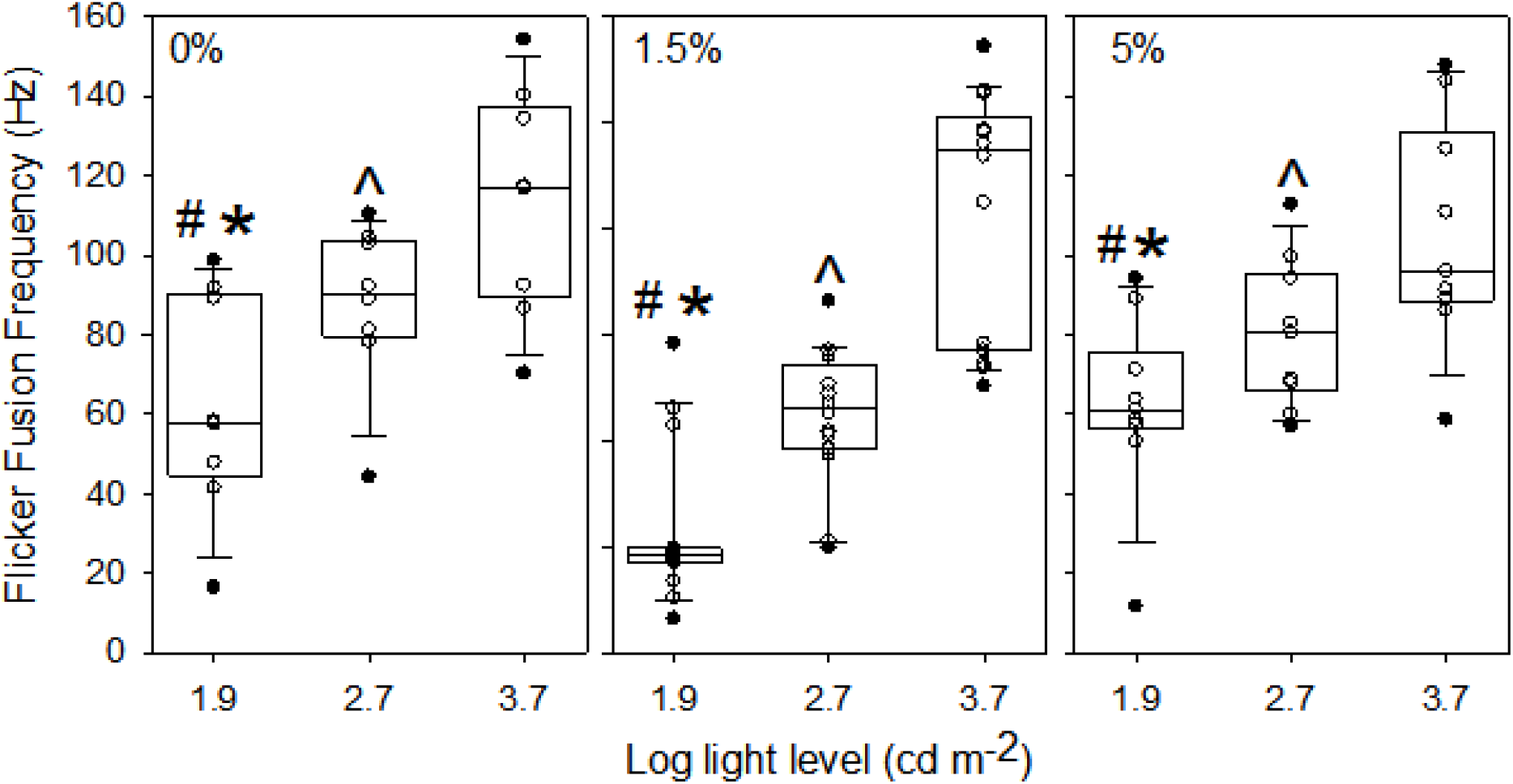
Flicker fusion frequencies (i.e., the highest frequency of sinusoidal light stimulus detectable) with increasing light intensities of European seabass. The light levels used correspond to approximately those needed to produce responses approximately 25%, 50% and 75% of the maximum response in the intensity–response curves (Fig 3). The symbols #, *, and ^ indicate differences in flicker fusion frequencies between log light levels 1.9 and 2.7, between log light levels 1.9 and 3.7, and between log light levels 2.7 and 3.7, respectively. The boundary of the box closest to zero indicates the 25^th^ percentile, the line within the box marks the median, and the boundary of the box farthest from zero indicates the 75^th^ percentile. Whiskers (error bars) above and below the box indicate the 90^th^ and 10^th^ percentiles. Data points above and below the 90^th^ and 10^th^ percentiles are considered outliers and are not used to determine median values.

The spectral sensitivity of fish fed diets with reduced taurine levels (i.e., 0 and 1.5%) were not significantly different from each other (Fig 6, panels A and B) and these data were subsequently combined. The results from fitting the data to both SSH and GFRKD rhodopsin templates suggested that European seabass are trichromats (i.e., have three visual pigments) (Tables 3 and 4). The SSH rhodopsin template was clearly best fitting (i.e., had the lowest AIC value) for the combined data from fish fed diets with reduced taurine levels (Table 3). In the case of data from fish fed 5% taurine diet (Table 4), the AIC values (i.e., goodness of fits) for SSH rhodopsin templates for dichromats (i.e., two retinal pigments) with secondary absorbing peaks (i.e., β bands) on both the visual pigments, and the SSH and GFRKD rhodopsin templates for trichromats, were indistinguishable (ΔAIC < 5). The SSH template for a dichromat with secondary absorbing peaks on both pigments predicted, however, the short wavelength pigment to have maximum absorbance in the UV wavelength (376 nm) which we consider unreasonable given the results from ERG data fitted similarly from a variety of inshore fishes (34-35, 41). We therefore based our subsequent conclusions on the effects diet on SSH rhodopsin templates for trichromats.

**Fig 6.**
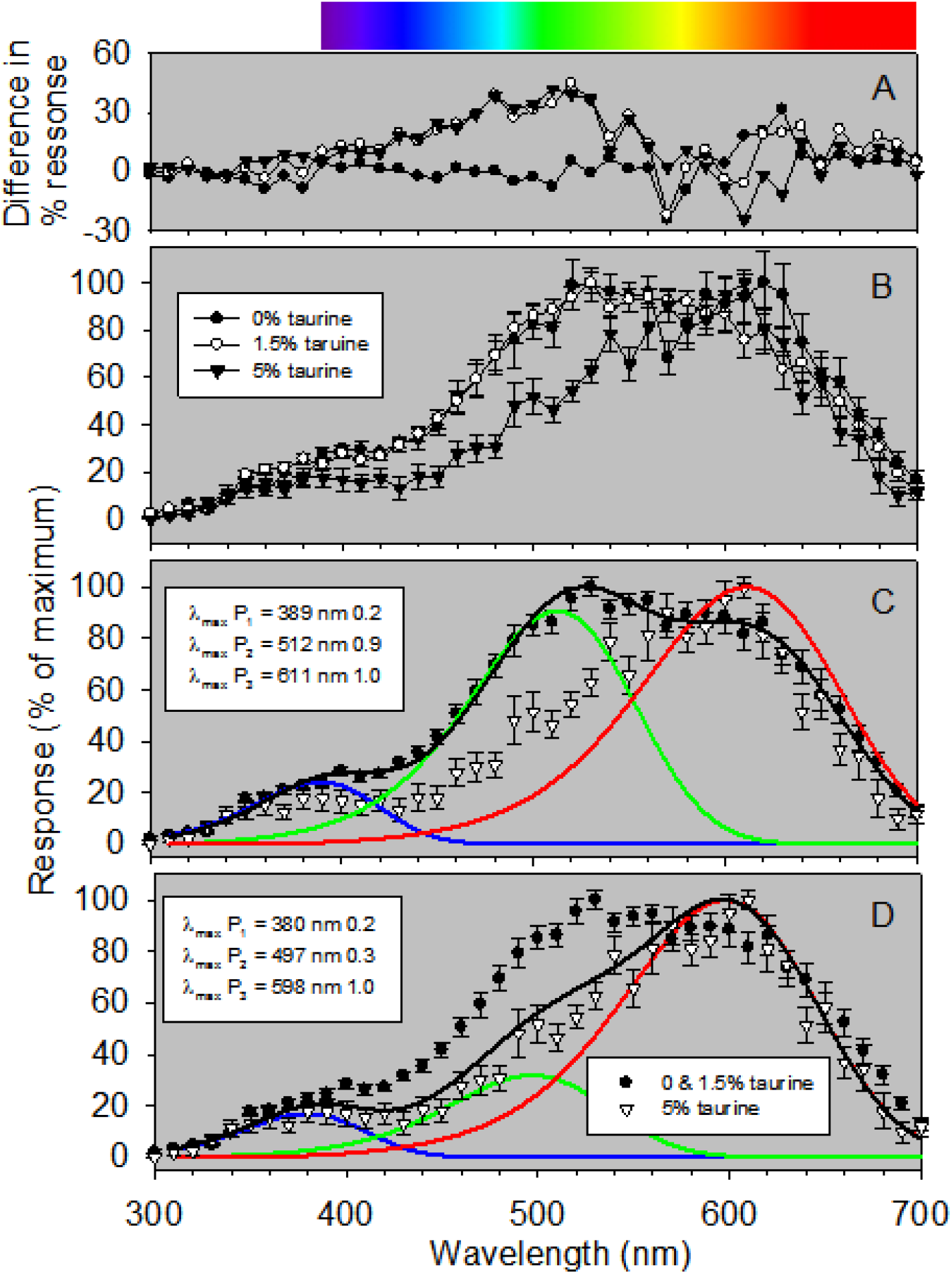
Spectral sensitivity (i.e., responses to monochromatic light) of European sea bass. Data are mean values (± SEM). Response values for individuals were normalized to 0-100%, the values averaged, and the mean values rescaled to 0-100%. Panel A are the differences between mean values of responses of fish fed the three different diets. Panel B are the mean responses of fish feed diets containing 0%, 1.5% and 5% taurine. In both panels, fish feed diets containing 0%, 1.5% and 5% taurine are indicated by filled circles, open circles and filled triangles, respectively. Because the spectral sensitivities of fish fed diets containing 0% or 1.5% taurine were largely not different, the data from these individuals were combined and the mean relative responses shown in panels B and C. Wavelengths at maximal absorptions (λ_max_ values) and pigment-specific weight values (to the right of each λ_max_ value) are shown. Black lines are the summed curves of the visual pigment curves multiplied by their respective weighting factors. Color coded lines indicate the absorptive characteristics of individual visual pigments. The color bar at the top of the figure shows the approximate human visual spectrum.

**Table 3.** The spectral sensitivity of European seabass fed diets with reduced taurine levels (i.e., 0 and 1.5%) where the data from fish in these two groups have been combined.

| Condition | Template | $\lambda_{\max,1}$ | $\lambda_{\max,2}$ | $\lambda_{\max,3}$ | $-\log(L)$ | p | AIC | $\Delta AIC$ | Pigment weights |
| --- | --- | --- | --- | --- | --- | --- | --- | --- | --- |
| Mono | GFRKD | 574 |  |  | -32.1 | 8 | -58 | 169 |  |
|  | SSH | 575 |  |  | -26.5 | 8 | -47 | 180 |  |
| Di, $\alpha$ | GFRKD | 410 | 510 | | -91.3 | 8 | -172 | 55 | |
|  | SSH | 411 | 510 |  | -91.0 | 8 | -173 | 55 |  |
| Di, $\beta$, S | GFRKD | 498 | 603 | | -79.7 | 8 | -147 | 80 | |
|  | SSH | 504 | 604 |  | -97.7 | 8 | -183 | 44 |  |
| Di, $\beta$, L | GFRKD | 439 | 513 | | -79.5 | 8 | -147 | 80 | |
|  | SSH | 461 | 513 |  | -96.5 | 8 | -181 | 46 |  |
| Di, $\beta$, B | GFRKD | 509 | 601 | | -67.2 | 8 | -120 | 107 | |
|  | SSH | 527 | 605 |  | -82.0 | 8 | -150 | 77 |  |
| Tri, $\alpha$ | GFRKD | 388 | 501 | 603 | -97.4 | 8 | -181 | 47 | |
|  | SSH | 389 | 512 | 611 | 120.7 | 8 | -227 | 0 | 0.24, 0.91, 1 |

**Table 4.** The spectral sensitivity of European seabass fed a 5% taurine diet.

| Condition | Template | $\lambda_{\max,1}$ | $\lambda_{\max,2}$ | $\lambda_{\max,3}$ | $-\log(L)$ | p | AIC | $\Delta AIC$ | Pigment weights |
| --- | --- | --- | --- | --- | --- | --- | --- | --- | --- |
| Mono | GFRKD | 586 |  |  | -52.2 | 8 | -98 | 92 |  |
|  | SSH | 588 |  |  | -44.6 | 8 | -83 | 108 |  |
| Di, $\alpha$ | GFRKD | 485 | 597 | | -83.7 | 8 | -155 | 35 | |
|  | SSH | 496 | 598 |  | -96.9 | 8 | -182 | 9 |  |
| Di, $\beta$, S | GFRKD | 485 | 597 | | -83.7 | 8 | -155 | 35 | |
|  | SSH | 496 | 598 |  | -96.9 | 8 | -182 | 9 |  |
| Di, $\beta$, L | GFRKD | 439 | 513 | | -79.5 | 8 | -147 | 44 | |
|  | SSH | 461 | 513 |  | -96.5 | 8 | -181 | 10 |  |
| Di, $\beta$, B | GFRKD | 506 | 597 | | -76.6 | 8 | -139 | 52 | |
|  | SSH | 376 | 599 |  | -102.4 | 8 | -191 | 0 |  |
| Tri, $\alpha$ | GFRKD | 382 | 492 | 597 | -101.4 | 8 | -189 | 2 | |
|  | SSH | 380 | 497 | 598 | -101.3 | 8 | -189 | 2 | 0.17, 0.32, 1 |

Our results suggested that the visual pigments of European seabass fed diets containing 0 or 1.4% taurine have maximal absorbance peaks (λ_max_) of 389, 512, 611 nm (P_1_, P_2_, P_3_, respectively; Fig 6, panel C). The maximal absorbance peaks of the visual pigments in fish fed a diet containing 5% taurine had maximal absorbance peaks (λ_max_) of 380, 497, 598 nm (P_1_, P_2_, P_3_, respectively; Fig 6, panel D). The most significant differences were a significant effect of dietary taurine level on the relative abundance of the three visual pigments (Fig 6, panel C). More specifically, in fish fed 0 and 1.5% taurine diets, the ratio of the middle and long wavelength visual pigments (P_2_ and P_3_, respectively) were approximately equal (0.91 to 1,) with a reduced abundance of the short wavelength pigment (P_1_) relative to the most abundant longwave length pigment (P_3_) (0.24 to 1) (Fig 6, panel C). By contrast, in fish fed a 5% taurine diet, there was a reduction in the abundance of the middle wavelength pigment (P_2_) relative to the long wavelength pigment (P_3_) (0.32 to 1) (Fig 6, panel D), while the ratio of the abundance of the short wavelength pigment (P_1_) relative to the most abundant longwave length pigment (P_3_) was relatively unchanged (0.17 to 1) (Fig 6, panel D). The overall effects of the changes in abundance of visual pigments was that the peak in overall retinal spectral sensitivity was shifted to longer wavelengths, from ≈530 nm in fish fed the 0% or 1.5% taurine diets (Fig 6, panel C), to ≈600 nm in fish fed the 5% taurine diet (Fig 6, panel D).

Replacing fishmeal with plant-based diets is a priority for the aquaculture industry, and the development of optimized diets is underway [10,42]. Plant-based diets have a high protein content [43], but commercial feed formulations often vary based on the availability and cost of ingredients, with different batches containing significantly different proportions or levels of quality of ingredients, or different ingredients all together. For this reason, the assessment of multiple ingredients and ingredient combinations is necessary. To address this issue, we formulated a diet based on available and cost-effective plant ingredients which are effective fishmeal replacements in rainbow trout (*Oncorhyncus mykiss*) [44–45] and are highly digestible by European sea bass [46]. But, as with other plant-based diets, our formulation lacks taurine. There is evidence that, although that taurine can be recycled through the taurine transporter and biliary recycling pathways, constant dietary supply is required to maintain proper function throughout multiple tissue types [20]. We assumed that taurine’s role as a photoreceptor protectant would most likely be as an antioxidant, similar to the role it plays in the liver of Atlantic salmon challenged with pro-oxidants like cadmium chloride [21,47]. We therefore hypothesized that visual disparities between European seabass fed diets containing 0%, 1.5% and 5% taurine would be due to differences in photoreceptor and pigment repair.

We did not, however, see a reduction of luminous sensitivity resulting from reduced dietary taurine (Fig 2). In contrast, other investigators have demonstrated a large progressive decrease in ERG amplitudes to a range of light intensities (12-197 cd m^-2^, log light level 1.1 to 2.3) in rats fed a soybean-based diet (containing a negligible amount of taurine) and treated with guanidinoethyl sulfonate (a compound that depletes of taurine) over 15 weeks [23]. Similar decreases in ERG amplitudes have been recorded in cats fed a taurine deficient (casein-based) diet (40). These results (in comparison to ours) imply a loss of luminous sensitivity resulting from a low taurine diet is related to a reduction in the cellular defense mechanisms countering light- and oxygen induced damage in retinal receptor cells [24], and that taurine deficiency has different effects in mammals and teleost fishes.

Effective visual function at low light levels requires both spatial and temporal summation within the retina [50–51]. The former results from the convergence of photoreceptors onto bipolar cells, as seen in fishes occupying dimly lit environments [52–54]. Spatial summation therefore reduces visual acuity (i.e., the ability to detect the details of an object). Demonstrating anatomical changes in spatial summation (i.e., changes in the convergence of photoreceptors onto bipolar cells) was, however beyond the scope of this study. Temporal summation refers to photoreceptors responding slowly to flashing light as evinced by temporal resolutions (i.e., reducsed FFF). Low FFF implies a blurring of fast-moving objects or the inability to detect them at all [50–51, 55–57]. We did specifically investigate the effects of dietary taurine level on flicker fusion frequency (Fig 5) but could we could not demonstrate any effects of dietary taurine level. We therefore conclude that there was no effect of dietary taurine level on retinal anatomy in terms of the degree of the convergence of receptor cells onto bipolar cells.

In terms of spectral sensitivity, our results imply that the European seabass visual system has three visual pigments and, more importantly, that spectral sensitivity can be impacted by diet taurine content (Fig 6). To the best of our knowledge, we are the first to demonstrate this phenomenon. Our results could be due to: (1) changes in the relative abundance of the three visual pigments equivalent to the seasonal shifts seen in other teleost fishes [58–60], or (2) changes in the amino acid composition of opsin proteins (which influence peak wavelength absorbancies of visual pigments) that result from changes in amino acid composition of opsin proteins. Changes in amino acid composition of opsin proteins (due to changes in gene expression) have been shown to occur over ontogeny, through adaptation of individuals to different light spectra, or changes diurnal lighting patterns [61–67]. Our data do not allow us to differentiate between changes in the relative abundance of visual pigments or changes in amino acid composition of opsin proteins. Our results do, however, demonstrate that there is a threshold effect of dietary taurine in European seabass, in that there were no discernable alterations in spectral sensitivity of fish fed 0% and 1.5% taurine diets, but a clear difference spectral sensitivity of fish fed a 5% taurine diet. The net effect of reduced taurine diets was a broader overall spectral sensitivity, with the peak in spectral sensitivity being leftward-shifted (a greater sensitivity to shorter wave lengths).

We note, however, that one of reasons for undertaking this study was to determine if fishes being cultured for stock enhancement programs fed a plant-based diet lacking sufficient taurine could decrease survival and fitness following release. If so, programs rearing fish for restocking would be less able to meet their ultimate objective of enhancing population abundance. Our demonstration that reduced dietary taurine levels influence spectral sensitivity (Fig 6) implies that this is the case. This conclusion is congruent with both the visual pigment sensitivity hypothesis and the contrast sensitivity hypothesis. The former posits that for optimal visual function the absorbance of visual pigments must correspond to the spectral distribution of the light environment [68–70], whereas the latter contends that aquatic animals maximize contrast sensitivity to acquire information effectively in a turbid (i.e., low contrast) environment [71–73]. Under both situations, however, changes in the relative abundance of visual pigment is likely to reduce fitness in cultured fish released for restocking. It unknown, however, if these deficits would remain following release of cultured, fish or if the visual properties of fish fed 0% and 1.5% taurine diets we observed would converge onto those of fish fed a 5% taurine diet after the switch to a natural prey diet. These questions are worth investigating.

## Conclusions

We could not demonstrate any effects of dietary taurine level on retinal anatomy, the functional properties of luminous sensitivity nor temporal resolution (i.e., FFF) in European sea bass. We did, however, find an effect on spectral sensitivity. The peak of spectral sensitivity of individuals fed a 5% taurine diet was rightward shifted (i.e., towards longer wavelengths) relative to that of fish fed a 0% or 1.5 % taurine diet. This difference in spectral sensitivity was caused by a relatively lower level of middle wavelength pigment (maximum absorbance ≈500 nm) in fish fed a 5% taurine diet. Changes in spectral sensitivity resulting from diets containing different taurine levels are unlikely to be detrimental to fish destined for market, but could be in fishes that are being reared for stock enhancement programs.

## Acknowledgements

The authors would like to thank the staff of the Aquaculture Research Center at the Institute of Marine and Environmental Technology, including Steve Rodgers and Chris Tollini. This is contribution XXXX from the Virginia Institute of Marine Science. This publication was made possible by the National Oceanic and Atmospheric Administration, Office of Education Educational Partnership Program award number (NA11SEC4810002). Its contents are solely the responsibility of the award recipient and do not necessarily represent the official views of the U.S. Department of Commerce, National Oceanic and Atmospheric Administration. The views expressed in this article are those of the authors and may not reflect the official policy of the Department of Health and Human Services, the U.S. Food and Drug Administration, or the U.S. Government.

